# Feline leukaemia virus subgrouping using the viral interference assay

**DOI:** 10.1101/2025.07.21.665799

**Authors:** Dimas Arya Abdillah, Takahiro Hiratsuka, Fumiko Matsuyama, Yuki Hattori, Takehisa Soma, Tetsuya Shimoda, Takashi Kato, Hideo Sakai, Masaharu Hisasue, Ariko Miyake, Kazuo Nishigaki

## Abstract

Feline leukaemia virus subgroup A (FeLV-A) is transmitted among cats, and FeLV subgroups are frequently generated *de novo*. We investigated the frequency of detection of subgroups using interference assays in 50 cases. FeLV-A infection alone was detected in 38% of cases, whereas co-infection with both FeLV-A and FeLV-B was observed in 62% of cases. No cases of FeLV-B infection alone were observed. Cases of co-infection with FeLV-A and FeLV-B showed a higher prevalence than cases of FeLV-A infection alone. The X region containing FeLV was discovered in two new cases. This study may elucidate the mechanism underlying FeLV-B-induced diseases.

---

Feline leukaemia virus (FeLV) was identified as the infectious agent responsible for feline leukaemia and lymphoma in the early 1960s [1]. FeLV infections are common worldwide and lead to malignant haematopoietic disorders, including lymphoma, myelodysplastic syndrome, acute myeloid leukaemia, aplastic anaemia, and immunodeficiency in cats. Cats of different ages can be infected with FeLV; however, kittens are more susceptible to progressive FeLV infection than adult cats. Cats with FeLV infection shed a large number of particles, and the infection is often fatal [1].

FeLVs can be categorised into several subgroups based on viral receptor interference [2]. FeLV subgroups, such as FeLV-A, -B, -C, -D, -E, and -T, have been identified [2, 3, 4, 5]. FeLV-B arises from recombination in the *env* region between FeLV-A and endogenous FeLV present in the feline genome [6]. Epidemiological and interference tests have shown that FeLV-A is an exogenous virus transmitted among domestic cats [1]. FeLV-B occurs *de novo* in cats already infected with FeLV-A. FeLV-B cannot cause cell-free infections independently [7]. Horizontal transmission of FeLV-B does not occur; however, it may rarely be transmitted by co-infection with FeLV-A [8, 9]. FeLV-B is frequently identified in cats with lymphoma [10]. The occurrence of myeloid leukaemia has been reported in mixed infections with FeLV-A and FeLV-B [11]. However, the role of FeLV-B in FeLV infections remains largely unknown.

The FeLV p27 antigen is the most common method for detecting FeLV infection using enzyme-linked immunosorbent assay [1]. However, antigen detection and antibody tests have not yet been developed to identify viral subgroups. In some cases, polymerase chain reaction (PCR) may be effective in detecting recombinant viruses, including FeLV-B and FeLV-D [4, 10]. Classically, receptor interference assays have been used to identify FeLV subgroups using mouse sarcoma virus, alkaline phosphatase, β-galactosidase, and green fluorescent protein as markers [2, 3, 4, 12]. Receptor interference assays have been widely used to identify FeLV subgroups. This is based on the principle that a virus cannot infect a cell that is already occupied by another virus that uses the same entry receptor. Initially, researchers used mouse sarcoma virus-based systems to classify FeLV subgroups; however, these assays required highly skilled techniques to assess viral infectivity in cells. Subsequently, marker genes such as alkaline phosphatase and green fluorescent protein were introduced to enhance detection. Despite improved visualisation, these approaches still required specialised and expensive equipment, such as fluorescence microscopes.

In this study, an improved and accessible receptor interference assay was developed using FeLV Env-pseudotyped viruses containing the LacZ reporter gene. These pseudo-viruses can enter cells through subgroup-specific Env– receptor interactions. Using 5-bromo-4-chloro-3-indolyl-β-D-galactopyranoside (X-gal) staining, FeLV subgroup classification is simpler, more visual, and easier to quantify. This system allows the direct, functional identification of FeLV subgroups based on receptor interference patterns, providing a practical tool for research and diagnostic applications.

In our previous study, we identified a novel recombinant FeLV containing an X region derived from *Felis catus* endogenous gamma retrovirus 4 (FcERV-gamma4) [13]. This recombinant FeLV was named XR-FeLV and was found in 6.4% of the FeLV-infected cats examined. Additionally, XR-FeLV was identified in 12.5% of 32 FeLV-infected cats diagnosed with lymphoma. All the isolated XR-FeLV clones shared a common X region that incorporated the middle portion of the FcERV-gamma4 5′ leader region. This sequence, present in all X regions, is also found in at least 13 endogenous retroviruses across the genomes of cats, humans, primates, and pigs and has been termed the common shared sequence [13]. However, the role of XR-FeLV in FeLV infections remains largely unknown.

In this study, we investigated the pathogenicity of FeLV-B based on clinical findings. To achieve this, we refined the method for studying FeLV subgroup infections, including FeLV-A, -B, -C, and -E.

To investigate the frequency of FeLV subgroup infection in clinical samples, we used a previously developed virus interference method [9]. Samples from FeLV-positive cats were generously provided by Drs. Fumiko Matsuyama (Yatsushiro Pet Clinic), Tetsuya Shimoda (Sanyo Animal Hospital), Takashi Kato (Kato Veterinary Hospital), Hideo Sakai (Isahaya Pet Clinic), and Takehisa Soma (Veterinary Diagnostic Laboratory, Marupi Lifetech Co., Ltd.).

For virus isolation (Fig. 1a), feline specimens such as serum/plasma (100 µL) or blood (30 or 100 µL) were inoculated into feline AH927 cells [5] in a 24-well plate and incubated for 2 days. The cells were washed to remove the inoculation material from the culture system, and then AH927 cells were cultured for an additional 2 weeks. DNA was purified from the recovered cells, and the culture supernatant was stored at -80 °C. Alternatively, for FeLV-B preparation, the supernatants of AH927 cells inoculated with specimens were transfected into human embryonic kidney transformed with SV40 large T antigen (HEK293T) cells [5] or Madin-Darby Bovine Kidney (MDBK) cells. FeLV was detected using PCR with template DNA from AH927, HEK293T, and MDBK cells, from which the viruses were isolated. KOD ONE polymerase (Toyobo, Osaka, Japan) was used. The FeLV *gag* gene was detected using primers Fe-23S (5′-AGGGATCCCAGCAGAAGTTTCAAGGCCACT-3′) and Fe-48R (5′-TTGAATTCCYGTGGCTCCTTGCACC-3′). The full-length *env* gene was detected using Fe-9S (5′-GAGACCTCTAGCGGCGGCCTAC-3′) and Fe-7R (5′-GTCAACTGGGGAGCCTGGAGAC-3′). The FeLV-B *env* gene was detected using specific primer PRB1(5′-CTGTTCACTCCTCGACAACG-3′) [10] and Fe-7R (5′-GTCAACTGGGGAGCCTGGAGAC-3′).

**Fig. 1.**
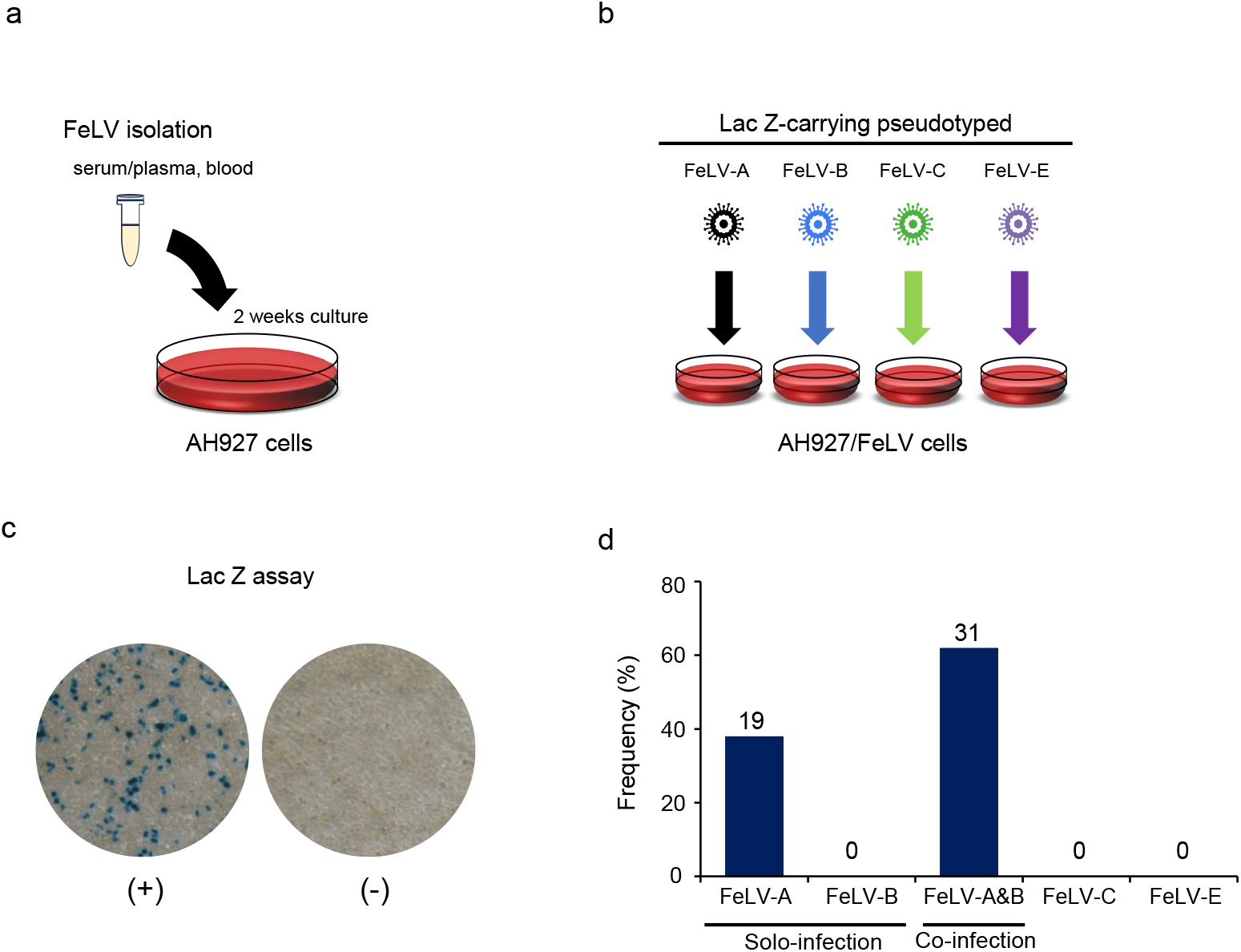
Schematic illustration of the viral interference assay. (a) Sample (serum/plasma or blood) was added to AH927 cells for 2 days, and the cells were then cultured for 2 weeks. (b) Subsequently, FeLV-infected AH927 cells were infected with each Lac Z-carrying pseudotyped virus for the classification of FeLV subgroups: FeLV-A, FeLV-B, FeLV-C, and FeLV-E. (c) Blue staining indicated galactosidase (LacZ)-positive cells, and blue-stained nuclei were counted under a microscope. (d) Virus interference assay was conducted in 50 cases. The results revealed the frequency of detection of FeLV subgroups as percentages (%). The numbers indicate the number of cases. FeLV, feline leukaemia virus

The preparation of Env-pseudotyped viruses carrying lacZ as a marker for the virus interference assay has been previously described [4]. GPLac cells were seeded at a concentration of approximately 1 × 10^6^ cells into a six-well plate 1 day before transfection. Expression vectors containing Env genes of FeLV-A (Glasgow-1), FeLV-B (Gardner–Arnstein), FeLV-C (Sarma), and FeLV-E (TG35-2) were transfected into GPLac cells (an env-negative packaging cell line containing the MLV *gag-pol* gene and a beta-galactosidase (LacZ)-coding pMXs retroviral vector) using TransIT-293 transfection reagent (Takara, Shiga, Japan), and cultured for 2 days. Culture supernatants were obtained as a source of FeLV subgroup of the Env-pseudotyped viruses carrying LacZ and were filtered through a 0.22-µm filter or centrifuged at 15,400 × *g* for 2 min at 4 °C, and then stored at -80 °C as virus stock for further experiments. For the virus interference assay (Fig. 1b), AH927 and FeLV-infected AH927 cells (approximately 3 × 10^5^ cells/well) were seeded into 24-well plates 1 day before infection. Target cells were infected with Env-pseudotyped virus (FeLV-A, FeLV-B, FeLV-C, and FeLV-E) (250 µL) in the presence of 10 µg/mL of polybrene (Nacalai Tesque, Kyoto, Japan) for 2 h. The cells were cultured for 2 days post-infection, and then, X-gal staining was conducted (Fig. 1c). The cell supernatants were removed, and the cells were fixed with 250 μL of 2% glutaraldehyde for 15 min at room temperature (20–25 °C), stained with X-gal, and counted as blue-stained nuclei under a microscope (Olympus CKX31, Hachioji, Tokyo). Viral titres are expressed as infectious units (IU) per millilitre (mL) with standard deviations. Typically, the viral titre in AH927 cells is approximately 1 × 10^4^ IU/mL. If a known subgroup of pseudoviruses infects FeLV-infected AH927 cells, it shows that superinfection has occurred, but viral interference has not. These results showed that the same subgroup of FeLV as the pseudovirus used in the assay is absent in the FeLV-infected AH927 cells. Conversely, if superinfection did not occur and viral interference was observed, it indicated that the same FeLV subgroup as the pseudovirus used in the assay was present in the FeLV-infected AH927 cells (Fig. 1c).

We successfully isolated FeLVs from 50 clinical samples using AH927 cells (Fig. 1d). A virus interference assay on these virus isolates revealed that 19 samples (38%) were positive for FeLV-A alone and 31 samples (62%) were positive for co-infection with FeLV-A and FeLV-B. None of the samples were infected with FeLV-B alone. No samples were infected with FeLV-C or FeLV-E. The presence of FeLV-B was confirmed using PCR. However, FeLV-B-specific PCR covering all samples has not yet been established; therefore, a cloning method for full-length Env was conducted using pCR4Blunt-TOPO (Invitrogen, Carlsbad, CA). FeLV-B was confirmed by sequencing at Fasmac Corporation (Atsugi, Japan).

Furthermore, PCR was conducted to investigate the presence of XR-FeLV in the samples from which the virus was isolated (Online Resource 1: Fig. S1a, S1b). The detection of XR-FeLV utilised Fe-23S (5′-AGGGATCCCAGCAGAAGTTTCAAGGCCACT-3′) and Fe-110R (5′-TGACTCCCAACAATCATTCCA-3′) primers. KOD FX Neo (TOYOBO, Osaka, Japan) polymerase was used, and the PCR amplicons were sequenced. XR-FeLV was found in all three samples. One of these samples was infected with FeLV-A only, whereas the other two were infected with both FeLV-A and FeLV-B (Online Resource 1: Table S1). Differences were observed in the recombination sites and lengths of the recombination regions of FeERV-gamma4 in each XR-FeLV X region (Online Resource 1: Fig. 2a, 2b, 2c). These results indicated that FeLV-B is frequently co-infected with FeLV-A in clinical samples. Subsequently, XR-FeLV was isolated from these cultures.

**Fig. 2.**
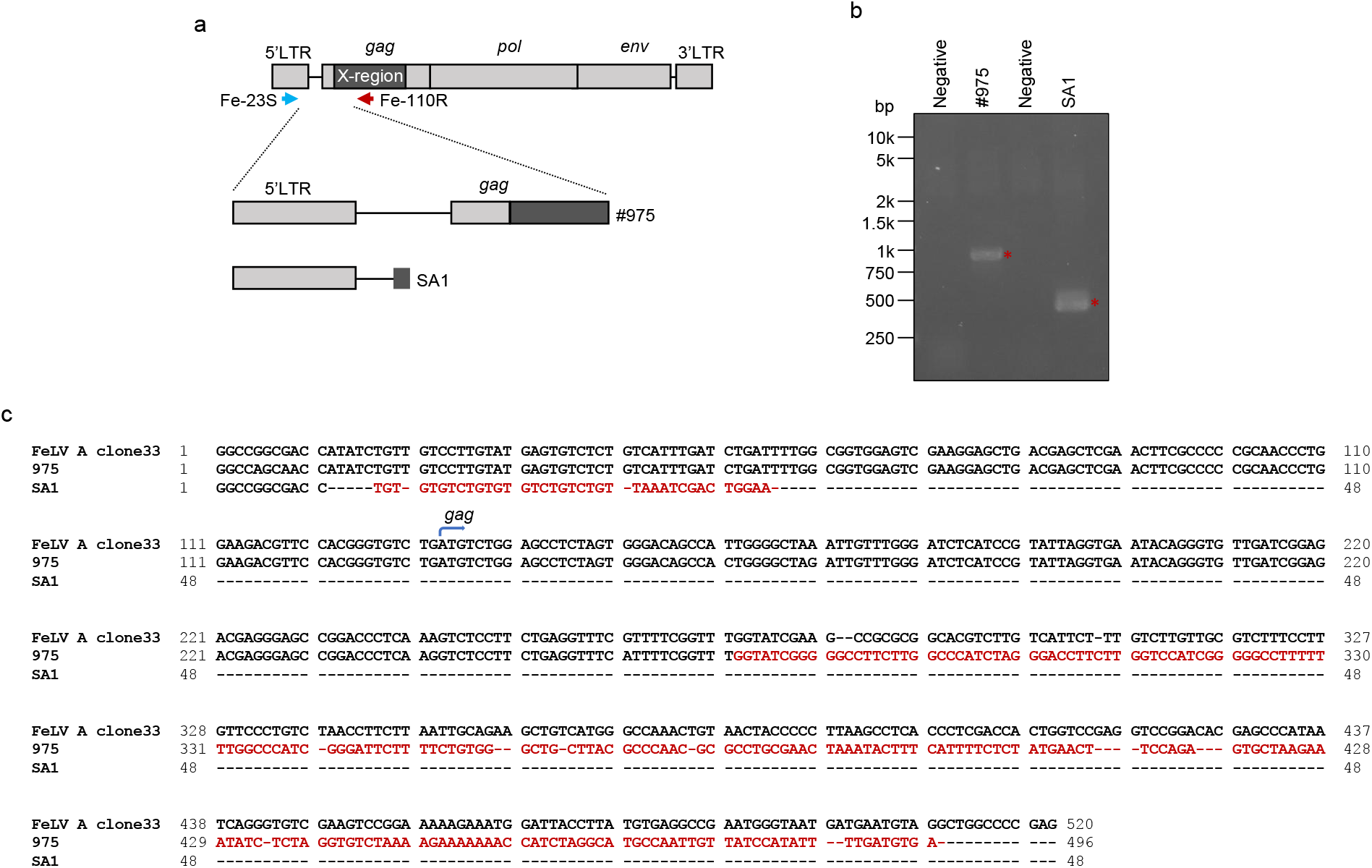
XR-FeLV detection using specific PCR primers. (a) Schematic representation of XR-FeLV obtained in this study. The black box indicates the X region derived from FcERV-gamma4, and the grey box indicates FeLV. Specific primers, Fe-23S and Fe-110R, used to detect XR-FeLV are indicated. Two samples (#975 and SA1) contained the X region. (b) Results of XR-FeLV detection using PCR. The asterisk indicates the specific bands of XR-FeLV. XR-FeLV, X region containing FeLV; FeLV, feline leukaemia virus; PCR, polymerase chain reaction. (c) Alignment of the nucleotide sequences of the X region compared with FeLV clone33. The red font indicates the X region of each sample.

Next, the relationship between FeLV subgroups and clinical findings in cats was examined (Table 1). There was no significant difference in age or sex between the FeLV-A solo-infected and FeLV-A/FeLV-B co-infected groups (age range: *P* > 0.05, sex ratio: *P* > 0.05) (Online Resource 1: Tables S2 and S3). Cases of co-infection with FeLV-A and FeLV-B (n=17) showed a higher prevalence than those with FeLV-A infection alone (n=17) (*p*<0.05 with Chi-squared test) (Table S4). When investigating the relationship between clinical symptoms and FeLV subgroups, we observed that FeLV-A infection was associated with anorexia and lymphoma (thoracic and submandibular). Contrastingly, cases of FeLV-A and FeLV-B co-infection presented with vomiting, stomatitis, renal failure, non-regenerative anaemia, immune-mediated haemolytic anaemia, and lymphoma (isolated renal and mediastinal types). There were no clear differences in clinical symptoms between the two groups, but when FeLV-B infection occurred, a tendency for systemic symptoms, including anaemia and renal disease, occurred.

**Table 1.**
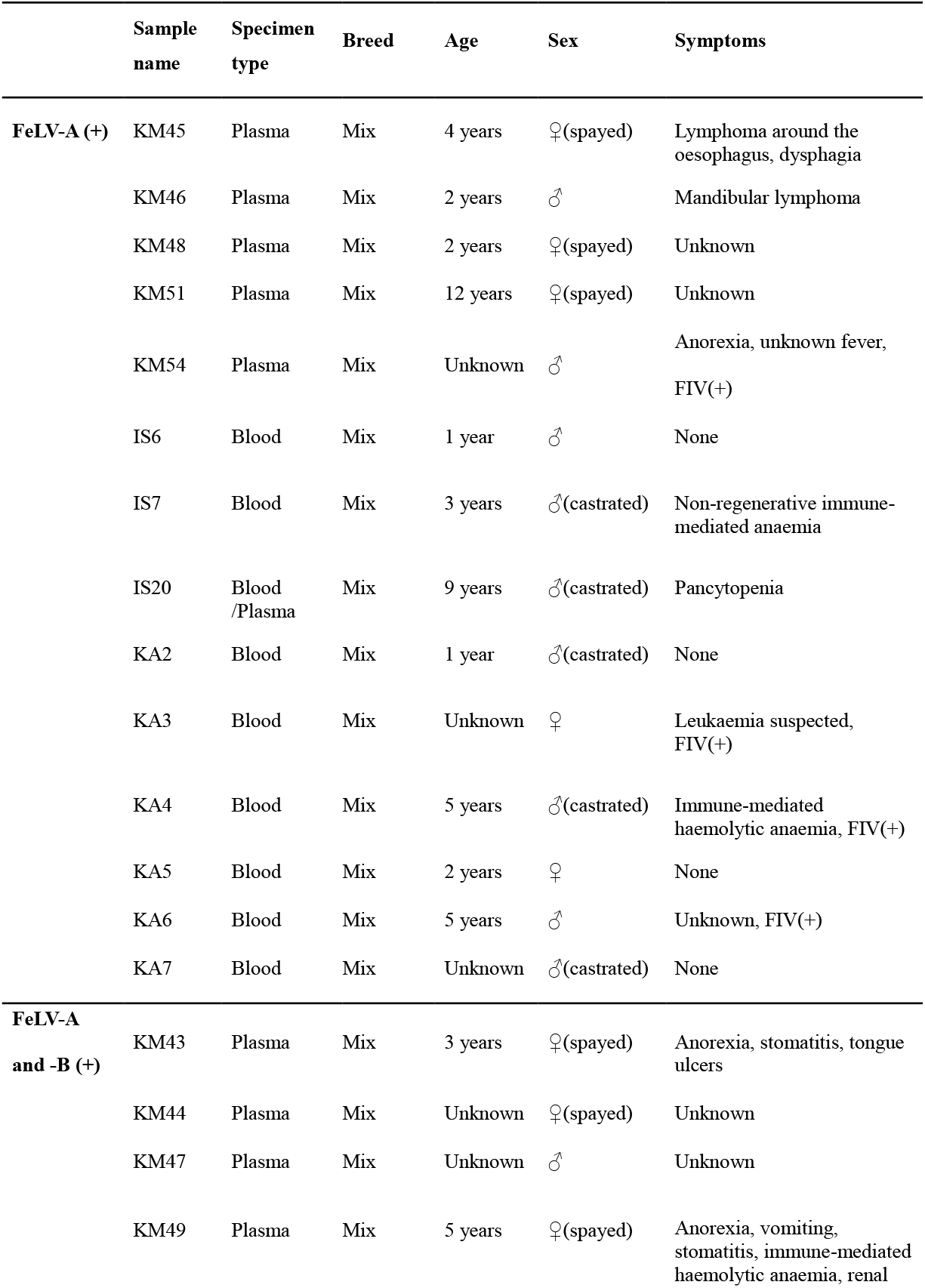
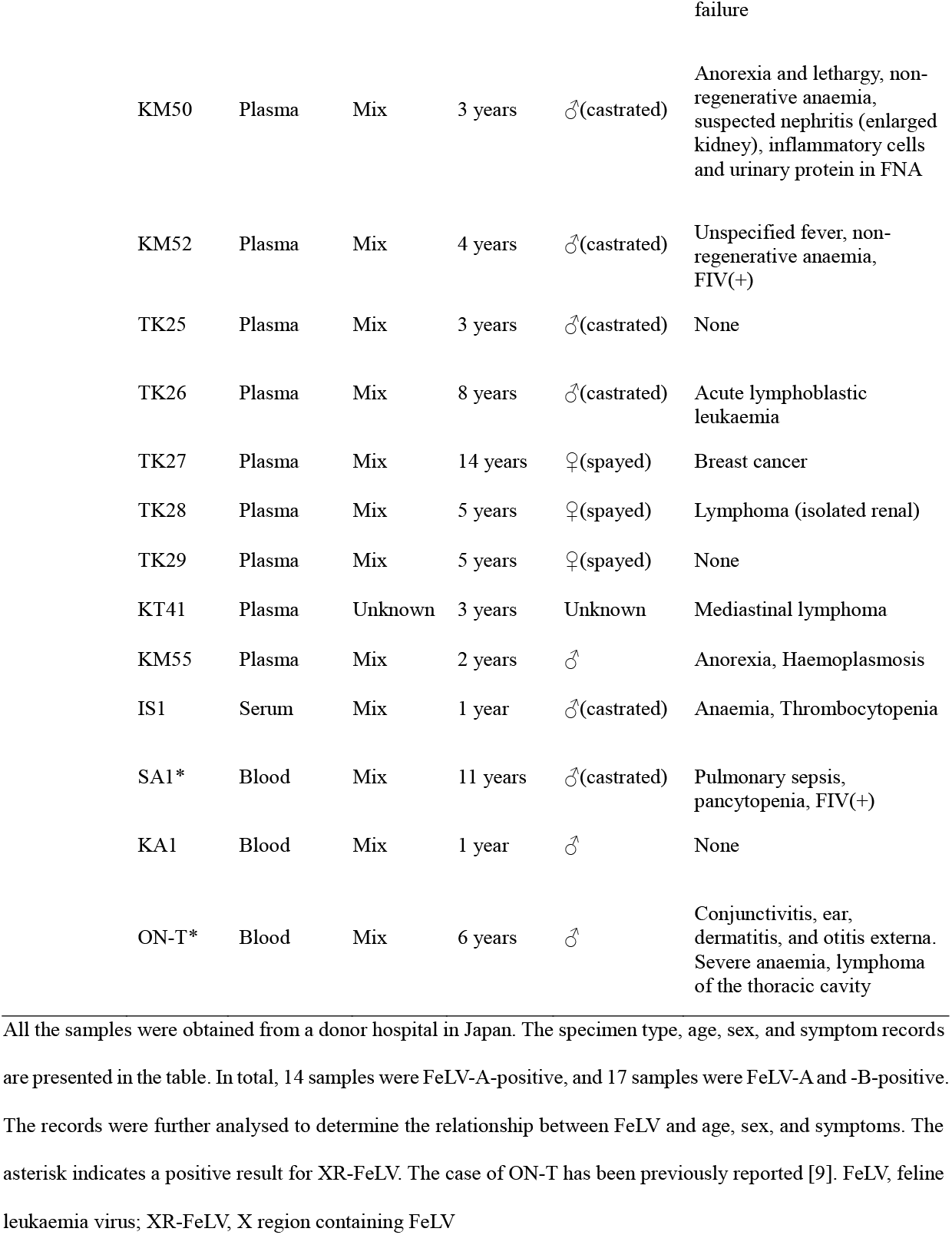
Clinical findings in FeLV-infected cats.

|  | Sample name | Specimen type | Breed | Age | Sex | Symptoms |
| --- | --- | --- | --- | --- | --- | --- |
| **FeLV-A (+)** | KM45 | Plasma | Mix | 4 years | ♀(spayed) | Lymphoma around the oesophagus, dysphagia |
|  | KM46 | Plasma | Mix | 2 years | ♂ | Mandibular lymphoma |
|  | KM48 | Plasma | Mix | 2 years | ♀(spayed) | Unknown |
|  | KM51 | Plasma | Mix | 12 years | ♀(spayed) | Unknown |
|  | KM54 | Plasma | Mix | Unknown | ♂ | Anorexia, unknown fever, FIV(+) |
|  | IS6 | Blood | Mix | 1 year | ♂ | None |
|  | IS7 | Blood | Mix | 3 years | ♂(castrated) | Non-regenerative immune-mediated anaemia |
|  | IS20 | Blood /Plasma | Mix | 9 years | ♂(castrated) | Pancytopenia |
|  | KA2 | Blood | Mix | 1 year | ♂(castrated) | None |
|  | KA3 | Blood | Mix | Unknown | ♀ | Leukaemia suspected, FIV(+) |
|  | KA4 | Blood | Mix | 5 years | ♂(castrated) | Immune-mediated haemolytic anaemia, FIV(+) |
|  | KA5 | Blood | Mix | 2 years | ♀ | None |
|  | KA6 | Blood | Mix | 5 years | ♂ | Unknown, FIV(+) |
|  | KA7 | Blood | Mix | Unknown | ♂(castrated) | None |
| **FeLV-A and -B (+)** | KM43 | Plasma | Mix | 3 years | ♀(spayed) | Anorexia, stomatitis, tongue ulcers |
|  | KM44 | Plasma | Mix | Unknown | ♀(spayed) | Unknown |
|  | KM47 | Plasma | Mix | Unknown | ♂ | Unknown |
|  | KM49 | Plasma | Mix | 5 years | ♀(spayed) | Anorexia, vomiting, stomatitis, immune-mediated haemolytic anaemia, renal |

|  |  |  |  |  |  |
| --- | --- | --- | --- | --- | --- |
|  |  |  |  |  | failure |
| KM50 | Plasma | Mix | 3 years | ♂(castrated) | Anorexia and lethargy, non-regenerative anaemia, suspected nephritis (enlarged kidney), inflammatory cells and urinary protein in FNA |
| KM52 | Plasma | Mix | 4 years | ♂(castrated) | Unspecified fever, non-regenerative anaemia, FIV(+) |
| TK25 | Plasma | Mix | 3 years | ♂(castrated) | None |
| TK26 | Plasma | Mix | 8 years | ♂(castrated) | Acute lymphoblastic leukaemia |
| TK27 | Plasma | Mix | 14 years | ♀(spayed) | Breast cancer |
| TK28 | Plasma | Mix | 5 years | ♀(spayed) | Lymphoma (isolated renal) |
| TK29 | Plasma | Mix | 5 years | ♀(spayed) | None |
| KT41 | Plasma | Unknown | 3 years | Unknown | Mediastinal lymphoma |
| KM55 | Plasma | Mix | 2 years | ♂ | Anorexia, Haemoplasmosis |
| IS1 | Serum | Mix | 1 year | ♂(castrated) | Anaemia, Thrombocytopenia |
| SA1* | Blood | Mix | 11 years | ♂(castrated) | Pulmonary sepsis, pancytopenia, FIV(+) |
| KA1 | Blood | Mix | 1 year | ♂ | None |
| ON-T* | Blood | Mix | 6 years | ♂ | Conjunctivitis, ear, dermatitis, and otitis externa. Severe anaemia, lymphoma of the thoracic cavity |

All the samples were obtained from a donor hospital in Japan. The specimen type, age, sex, and symptom records are presented in the table. In total, 14 samples were FeLV-A-positive, and 17 samples were FeLV-A and -B-positive. The records were further analysed to determine the relationship between FeLV and age, sex, and symptoms. The asterisk indicates a positive result for XR-FeLV. The case of ON-T has been previously reported [9]. FeLV, feline leukaemia virus; XR-FeLV, X region containing FeLV

Among the three samples from which XR-FeLV was isolated, only two showed clinically known symptoms. The symptoms of one sample (sample SA1) were pulmonary sepsis and pancytopenia, and those of the other sample (sample ON-T) were lymphoma, severe anaemia, conjunctivitis, ear dermatitis, and otitis externa [9]. In this study, a clear relationship between XR-FeLV and clinical symptoms could not be established owing to the small number of cases.

FeLV-A infection or co-infection with FeLV-A and FeLV-B is frequently observed in FeLV-infected cats. However, FeLV-B pathogenicity remains unknown because of the lack of *in vivo* experiments using FeLV-B alone in domestic cats. In this study, we investigated the pathogenicity of FeLV-B based on clinical findings. Infection with FeLV-B alone was not observed, and cases of co-infection with FeLV-A and FeLV-B showed a higher prevalence than cases of FeLV-A infection alone. It is believed that FeLV-B infection occurs after FeLV-A infection, leading to the pathogenesis of the FeLV infection complex.

In this study, we isolated FeLV field strains from clinical specimens using AH927 cells and established a series of systems for conducting interference experiments using a pseudotyped virus carrying the LacZ gene to confirm the FeLV subgroup. A new interference experimental system for FeLV-E was also included in this study. The *gag* gene was detected using PCR to confirm the successful isolation of FeLV. Enzyme-linked immunosorbent assay could also be used as an alternative method to identify the viral antigen. Feline-derived cells are considered ideal for virus isolation because FeLV-A replicates efficiently in feline cells. HEK293T cells were also tested for virus isolation; however, the efficiency was poor. FeLV isolated from AH927 cells could replicate in non-feline cells, such as 293T and MDBK cells used in this study. Once the virus has been successfully isolated from AH927 cells, it can efficiently replicate in non-feline cells owing to the high viral load. This may explain why FeLVs adapt to cultured cells. Sequence analysis revealed that these cell lines contained both FeLV-A and FeLV-B. As previously reported, there is large structural diversity in the FeLV-B *Env gene* sequence [10, 14]; therefore, the presence or absence of FeLV-B may not be accurately determined by identifying only recombinant viruses using PCR. FeLV-B-specific PCR was used in this study; however, it may be challenging to detect all FeLV-B strains. Thus, establishing a PCR method specific to FeLV-B is necessary.

Plasma, serum, or blood samples were used for virus isolation because FeLV-infected cats are viraemic. Another mention could involve co-culturing with peripheral blood mononuclear cells (PBMCs). However, obtaining PBMCs in clinical practice can be challenging. Virus isolation can be achieved using various biological materials, including serum, blood, organs, and saliva [1]. However, it should be noted that the isolation of subgroups may differ depending on the specific material used. Moreover, when the objective is to isolate unique FeLV strains, using non-feline and AH927 cells together for virus isolation could be effective. Whether FeLV-B is generated during virus isolation remains unknown. Therefore, developing a direct PCR method using biological samples is important.

The frequency of detection of viral subgroups was investigated in 50 viral isolates, with 100% of the cases showing FeLV-A infection and 62% of the cases showing a mixture of FeLV-A and FeLV-B infections. FeLV-A infection alone was observed in 38% of the cases, whereas FeLV-B infection alone was not observed. This result confirmed that FeLV-A was transmitted among cats during the FeLV infection. The fact that no single infection of FeLV-B was observed, despite 62% of infections containing an FeLV-B combination, suggests a potential resistance to FeLV-B infection. In close relationships, although rare, it is possible that FeLV-A and FeLV-B may be co-transmitted. However, the presence of FeLIX could prevent the transmission and generation of FeLV-B among cats through the FeLV-B receptor Pit1 [15]. In this study, there were no cases of FeLV-C or FeLV-E infection in the 50 virus isolates; therefore, conducting a large-scale survey would be required for more accurate information regarding these other strains.

The rate of FeLV infection is higher in young cats than in adult cats [14, 16, 17]. This phenomenon is hypothesised to be attributable to the fact that the immune system of young cats is still developing [16-18]. Furthermore, there is no difference in the rate of FeLV infection between the sexes [14]. This is because the main route of infection is not through fighting but through grooming and other activities. In this study, there was no significant difference in age or sex between the FeLV-A solo-infected and FeLV-A/FeLV-B co-infected groups (Online Resource 1: Tables S2 and S3). However, co-infection with FeLV-A and FeLV-B tended to have a higher prevalence than that of FeLV-A infection alone. Given that FeLV-B occurs *de novo* and appears after FeLV-A infection, there is a higher likelihood that FeLV-B will appear when the infection has progressed. When considering the mechanism of FeLV-B occurrence, the prevalence and FeLV-B infection rates may be correlated. FeLV-B infection has been frequently detected in lymphoma [10, 14]. Furthermore, the presence of FeLV-B may accelerate disease progression. While FeLV-A uses THTR1 for infection [19], FeLV-B is known to use Pit1 as the entry receptor [15]. Given that the target cells of viral infection differ, co-infection is expected to result in a wider range of target cells over a lifetime. The results of this study show that FeLV-B infection causes systemic symptoms and can be explained by the frequent detection of FeLV-B in tumours.

Given that FeLV-B infection occurs *de novo*, the frequency of replicating FeLV-B remains unknown. In this study, a virus isolated from AH927 cells was inoculated into cells (HEK293T and MDBK) intended for FeLV-B infection. Both FeLV-A and FeLV-B were detected in the cells. Since FeLV-A exists as a helper virus, the significance of FeLV-B as a replicating virus remains unknown.

Viral interference assay is a very powerful method for identifying known FeLV subgroups in FeLV isolates. Furthermore, this research method is also considered to be very useful for identifying unknown viral subgroups. However, if only FeLV-A is isolated in culture, it is unknown whether FeLV-B or other subgroups exist *in vivo*. Therefore, this technique hopefully develops into a method for identifying FeLV subgroups using genetic testing with biological materials.

It is unclear whether the presence (emergence) of XR-FeLV or subgroup FeLV other than FeLV-A contributes to the pathogenesis of FeLV infection. It is also possible that FeLV subgroups emerge as a result of disease progression. In a previous study, the clonality of FeLV-D was observed in a lymphoma case, suggesting that the emergence of viral subgroups is closely associated with tumorigenesis [9].

In this study, the relationship between the viral copy number and disease in cats was not examined. This will be a topic for future research.

In Japan, FeLV infection is a serious problem in some areas. Since the Japanese strain is genetically different from foreign strains [14], developing a vaccine using the Japanese strain to prevent FeLV infection may be necessary. Furthermore, in this study, we found a need to elucidate the pathogenesis of FeLV-A and FeLV-B. It is important to focus on both FeLV-A and FeLV-B as strategies for the prevention and treatment of FeLV infection in the future.

In this study, a method to identify FeLV subgroups was developed. FeLV infection is extremely complex owing to the emergence of different virus types after initial infection. The detection rate of FeLV-B infection in clinical specimens was relatively high, and the co-infection of FeLV-A and FeLV-B showed a higher prevalence. The results of this study show the importance of accurately evaluating the pathogenicity of FeLV-B. Further large-scale surveys will enable us to track the actual situation of the subgroups.

## Acknowledgments

This study was funded by the Japan Society for the Promotion of Science KAKENHI (grant number: 23K27086); K.N. received the funding. The funder had no role in the study design, data collection and interpretation, or the decision to submit the work for publication.

We are grateful to Hajime Tsujimoto for providing the FeLV-A/pFGA5 (Glasgow-1), FeLV-B/pFGB, and FeLV-C/pFSC plasmids and to Toshio Kitamura for the PLAT-GP cells and pMxs retroviral vector.

## Statements and Declarations

### Funding

This study was funded by the Japan Society for the Promotion of Science KAKENHI (grant numbers:23K27086); K.N. received the funding.

### Competing Interests

The authors declare no conflicts of interest.

### Author Contributions

The contributions of the authors are described as follows. Conceptualisation: K.N. Funding acquisition: K.N. Investigation: D.A.A., F.M., T.H., D.P., T.S., T.K., H.S., M.H., A.M., and K.N. Sources: F.M., Y.H., T.S., T.S., T.K., H.S., M.H., and K.N. Project administration: K.N. Supervision: K.N. Data analysis: D.A.A., T.H., A.M., K.N. Writing: D.A.A. and K.N.

## Data Availability

### Ethical Approval

Animal studies were conducted in accordance with the guidelines for the care and use of laboratory animals provided by the Ministry of Education, Culture, Sports, Science, and Technology, Japan. All the experiments were approved by the Genetic Modification Safety Committee of Yamaguchi University.

## Supplementary Information (SI): Online Resource 1

**Table S1.**
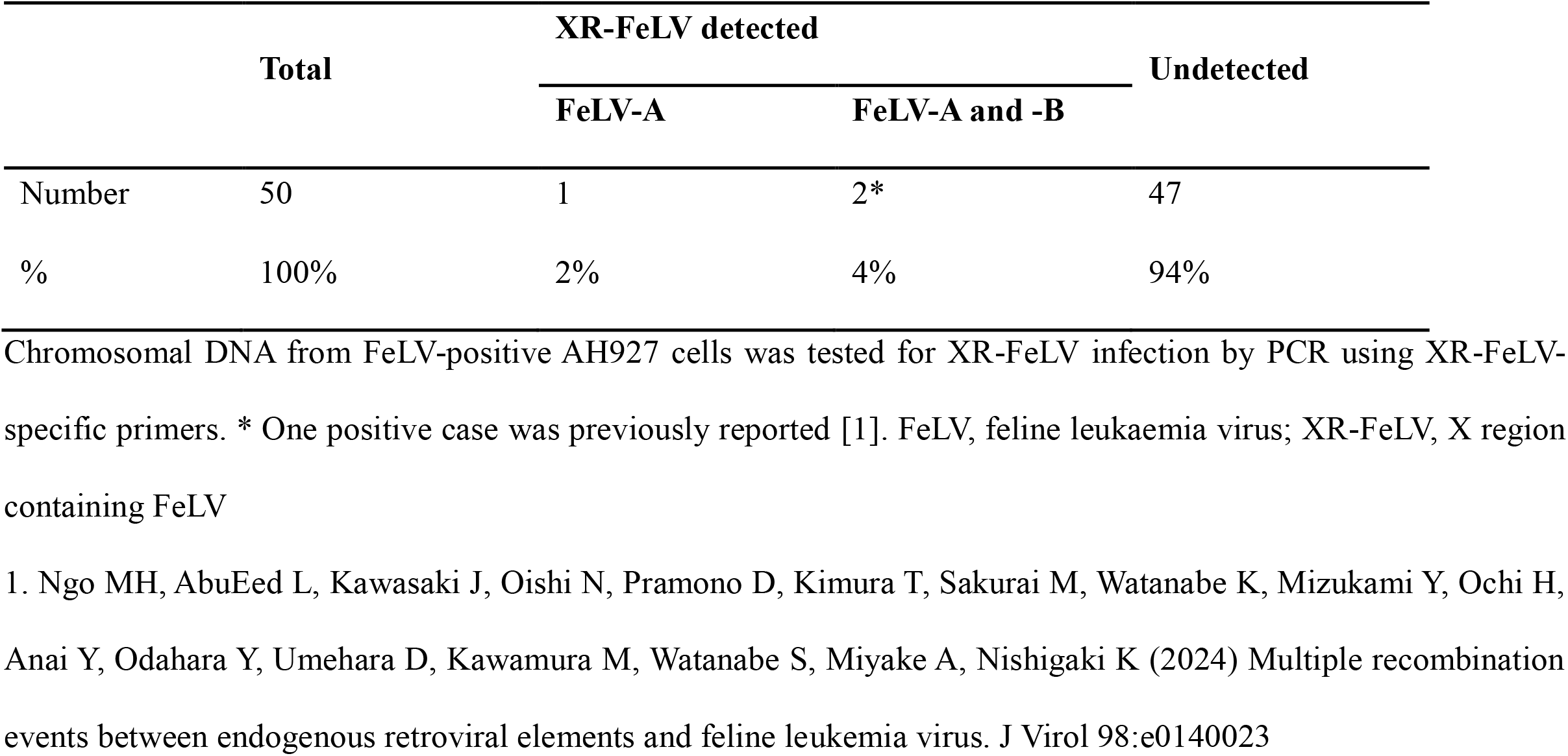
Frequency of XR-FeLV in FeLV isolates.

**Table S2.** Sex ratio.

| **Sex** | **FeLV-A** | **FeLV-A and B** |
| --- | --- | --- |
| ♂<sup>†</sup> | 9 | 10 |
| ♀<sup>#</sup> | 5 | 6 |

The sex ratio was calculated by comparing males and females in FeLV-A-infected cats (n=14) and FeLV-A and -B-infected cats (n=16). <sup>†</sup>Numbers including castrated. <sup>#</sup>Numbers including spayed. Unknown data were excluded from the analysis. The *P*-value for the sex ratio difference between FeLV-A and FeLV-A and -B was >0.05 (with no significant difference) with the Chi-squared test. FeLV, feline leukaemia virus

**Table S3.** Adult/Young ratio.

| **Age (year)** | **FeLV-A** | **FeLV-A and B** |
| --- | --- | --- |
| 1-6 [Adult] | 9 | 12 |
| >7 [Senior] | 2 | 3 |

The adult/senior ratio was calculated by comparing males and females in each FeLV-A (n=11) and FeLV-A and -B (n=15)-infected cat. Unknown data were excluded from the analysis. The *P*-value for the adult/senior ratio difference between FeLV-A and FeLV-A and -B was >0.05 (not a significant difference) with Chi-squared test. FeLV, feline leukaemia virus

**Table S4.**
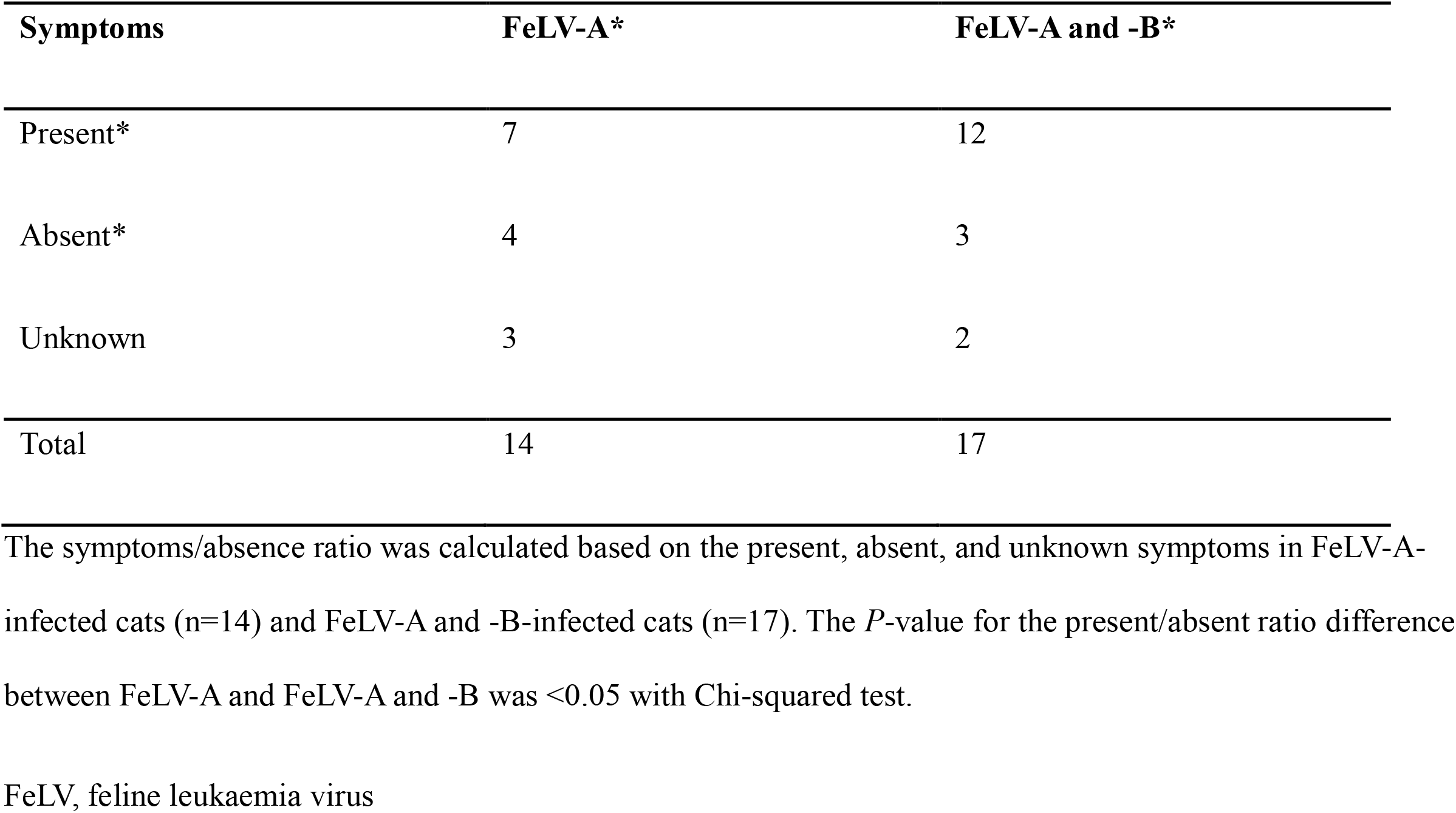
Symptoms present/absent ratio.

## Notes

### Competing Interest Statement

The authors have declared no competing interest.

